# Histolytics: A Panoptic Spatial Analysis Framework for Interpretable Histopathology

**DOI:** 10.1101/2025.08.26.672346

**Authors:** Oskari Lehtonen, Niko Nordlund, Shams Salloum, Ilkka Kalliala, Anni Virtanen, Sampsa Hautaniemi

## Abstract

Quantifying spatial organization in hematoxylin and eosin (H&E)–stained whole-slide images (WSIs) is essential for uncovering tissue-level patterns relevant to pathology. We present Histolytics, an open-source, scalable Python framework for interpretable, WSI-scale histopathological analysis. Histolytics integrates panoptic segmentation with spatial querying, morphological profiling, and graph-based analytics to enable high-resolution, quantitative characterization of nuclei, tissue compartments, and the extracellular matrix (ECM). Designed to align with diagnostic reasoning, Histolytics supports segmentation with state-of-the-art deep learning models and provides modular tools for extracting biologically grounded features across entire WSIs. By leveraging spatially contextualized measurements at cellular and tissue levels, Histolytics addresses a critical gap in explainable computational pathology, offering an interpretable alternative or complement to black-box predictive models. The framework is compatible with the broader Python data science ecosystem and includes extensive documentation and pretrained models to promote widespread adoption.

## Main

Hematoxylin and Eosin -stained (H&E) tissue Whole Slide Images (WSIs) form the cornerstone of histopathology. Their rich information content, coupled with their widespread availability, has presented a powerful resource for understanding diseases for centuries. Indeed, histopathology has a long and established history rooted in the visual assessment of features that are directly associated with various disease states, forming the basis of diagnosis and disease behavior. More recently, the advent of DL-approaches has introduced a powerful predictive modeling paradigm which has led to significant increases in computational prognostic accuracy, automated grading of tumors, and prediction of treatment response^1–5^. To increase the acceptance and trust in these AI predictions, improvements in textual and visual explainability have also been rapid^6–9^.

Despite the transformative potential of AI, the adoption of these models in routine histopathology workflows has been slow and applications remain mostly in the research settings^10^. A major limitation of these models is their inability to numerically quantify predefined or novel histology-informed features with biological relevance, such as tissue and cell type compositions, and nuclear and ECM morphology and organization. In contrast, recent methods utilizing segmentation and spatial analysis techniques have shown promise in such prognostics and diagnostics applications^11−13^. However, while significant efforts have been directed towards frameworks unifying DL-based histological predictive modeling under easy-to-use computational frameworks^14–16^, a similar effort to unify panoptic segmentation based spatial feature quantification methods under a comprehensive open-source framework is still lacking (Supplementary Table 1).

We developed Histolytics, an open-source Python framework designed to extract interpretable features that reflect human understanding and closely align with expert diagnostic reasoning. Histolytics uniquely combines full-slide panoptic segmentation^17^, which is a powerful approach to simultaneously segment tissues and nuclei, with advanced spatial analysis methods enabling interpretable, WSI-scale insights that reflect real-world tissue architecture and diagnostic reasoning. Histolytics also integrates with the wider Python ML ecosystem that together with modular application programming interface (API), detailed documentation, and extensive tutorials, ensure ease of use and integration into diverse computational pathology workflows. Histolytics is available at https://github.com/HautaniemiLab/histolytics with documentation and tutorials at https://hautaniemilab.github.io/histolytics/.

## Results

The design principle of Histolytics relies on three pillars, versatile WSI I/O, panoptic segmentation, and flexible spatial feature extraction (Fig 1). WSI I/O, a pre-requisite for panoptic segmentation and downstream spatial analyses at WSI-scale, is handled in Histolytics with a versatile WSI SlideReader-class. This class, through different slide-reading backends^18–20^, supports an extensive set of slide formats (Supplementary Table 2). Additionally, it includes slide-level masking and tile-filtering that enable the removal of WSI background and redundant sets of tissues (Fig 1a) for optimized panoptic segmentation run-times.

**Figure 1:**
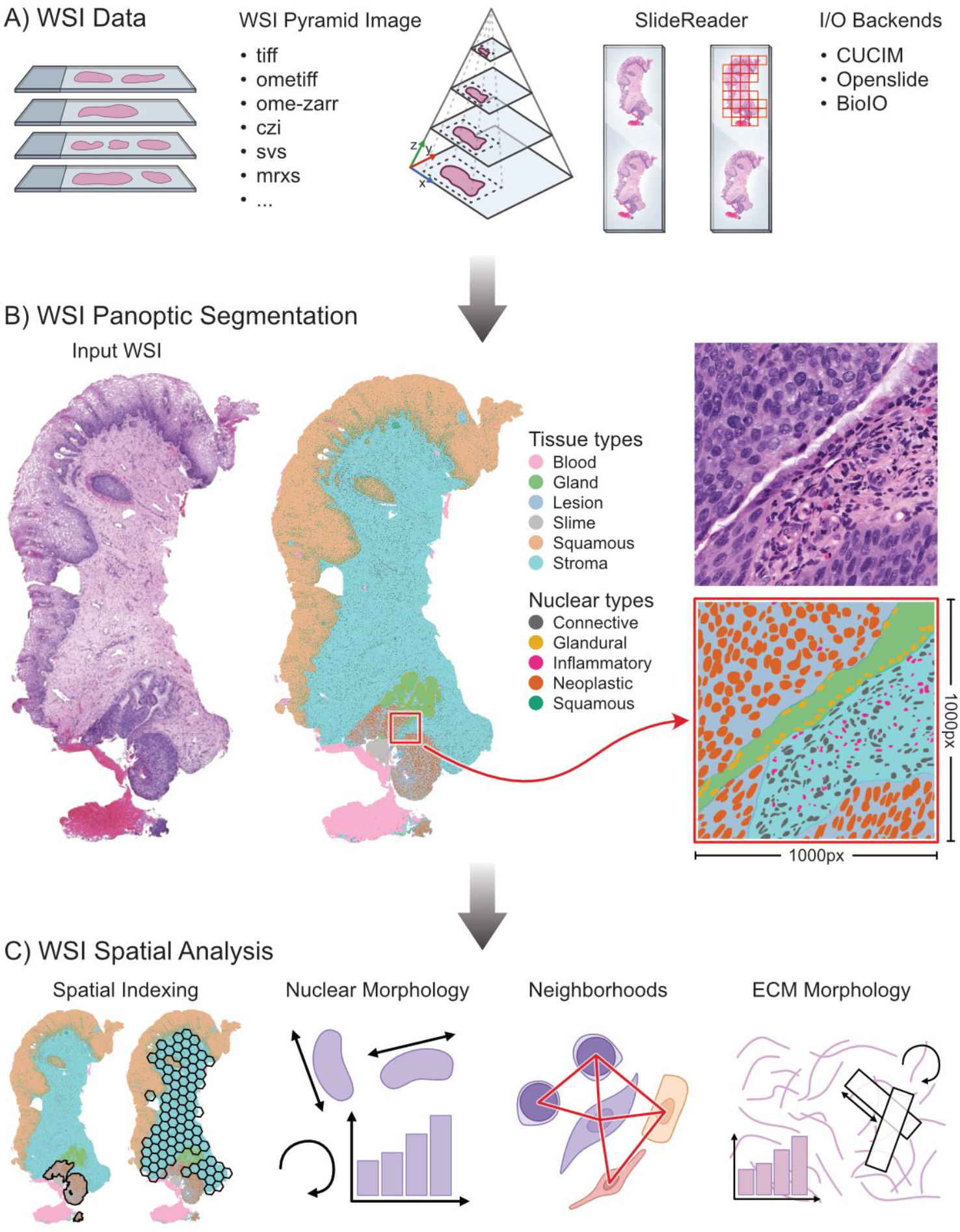
Histolytics is a spatial analysis library for histopathological images based on WSI-scale panoptic segmentation. **A)** Histolytics supports a wide range of WSI data formats through a versatile SlideReader-object with support for mask assisted tiling and a connected components-based tile selector. The SlideReader-object can be initialized with three different backends for WSI I/O: CUCIM, Openslide and BioIO. **B)** Illustration of WSI-scale panoptic segmentation maps, the primary engine of spatial analysis in Histolytics for spatially aware interpretable histopathological feature extraction. **C)** Histolytics spatial analysis is built upon a strong spatial indexing system and a wide array of methods for quantification of cell nuclei morphology, neighrborhood relationships, and extracellular matrix morphology.

To enable panoptic segmentation, Histolytics contains modular implementations of well-established cell segmentation models^21–24^ with tissue segmentation extensions. The modular implementations allow the easy integration of different model backbones such as the most recent histopathological foundation models^1–3^, allowing users to leverage the latest advancements in deep histological feature representations in their segmentation pipelines. To streamline fine-tuning and deployment, Histolytics offers several pre-trained models (available at https://huggingface.co/histolytics-hub) and a range of training and model validation tools, including multitask losses, regularization techniques, and performance evaluation metrics (Supplementary Table 2). Importantly, Histolytics includes tools to stitch patch-level segmentations into WSI-scale panoptic segmentation maps (Fig 1b), incorporating large-scale, comprehensive spatial context for every segmented object while erasing the need to operate at a tile-level.

The spatial analysis tools of Histolytics include efficient spatial querying and indexing for locating objects of interest and a wide range of morphological, textural, and graph-based feature extraction methods designed to extract interpretable histopathological features (Fig 1c). Built upon Libpysal^25^, Geopandas^26^, and a vectorized segmentation data format (GeoJSON, GeoParquet), Histolytics is optimized for large-scale geospatial analyses, with the familiar pandas DataFrame API^27^. This ensures a shallow learning curve and seamless integration with the broader Python data science and machine learning ecosystem while also exposing the powerful geospatial visualization engine of Geopandas. This also enables the usage of widely used image viewers like QuPath^28^ for interactive visualization of segmentation and spatial analysis outputs.

### WSI-Scale Cellular Organization Analysis via Spatial Querying

The quantification of the spatial organization of different cell types within different tissue compartments is fundamental in deciphering biological processes, particularly in complex environments like the tumor microenvironment^29,30^. For instance, to quantify Tumor Infiltrating Lymphocytes (TILs), a prognostic biomarker in several cancers^31–33^, the cancer and immune cell populations need to be precisely pinpointed within regions of interest. Mapping and counting specific nuclei within regions of interest can be challenging due to the potentially vast number of nuclei present in WSIs. Histolytics is designed to address this through efficient spatial querying and spatial partitioning tools applied to panoptic segmentation maps.

To demonstrate how Histolytics can be used for quantifying immune cell infiltration at WSI-scale, we used Histolytics’ cervical panoptic segmentation model and spatial querying tools to compare the levels of immune infiltration in two cervical intraepithelial lesions, a precursor to cervical cancer that is caused by a viral infection which can provoke host immune responses^34^. To add granularity to the analysis, we also used Histolytics’ interface-region tool to partition the segmented stromal tissues into two distinct sectors relative to the lesions: an adjacent zone representing the lesion-stroma interface and the distal stromal region (Fig 2a). This allowed us to compare the immune cell counts and proportions within the lesions, at the interface area, and at the distal stromal regions (Fig 2b, Supplementary Fig 1 shows group-level comparisons at cohort level).

**Figure 2:**
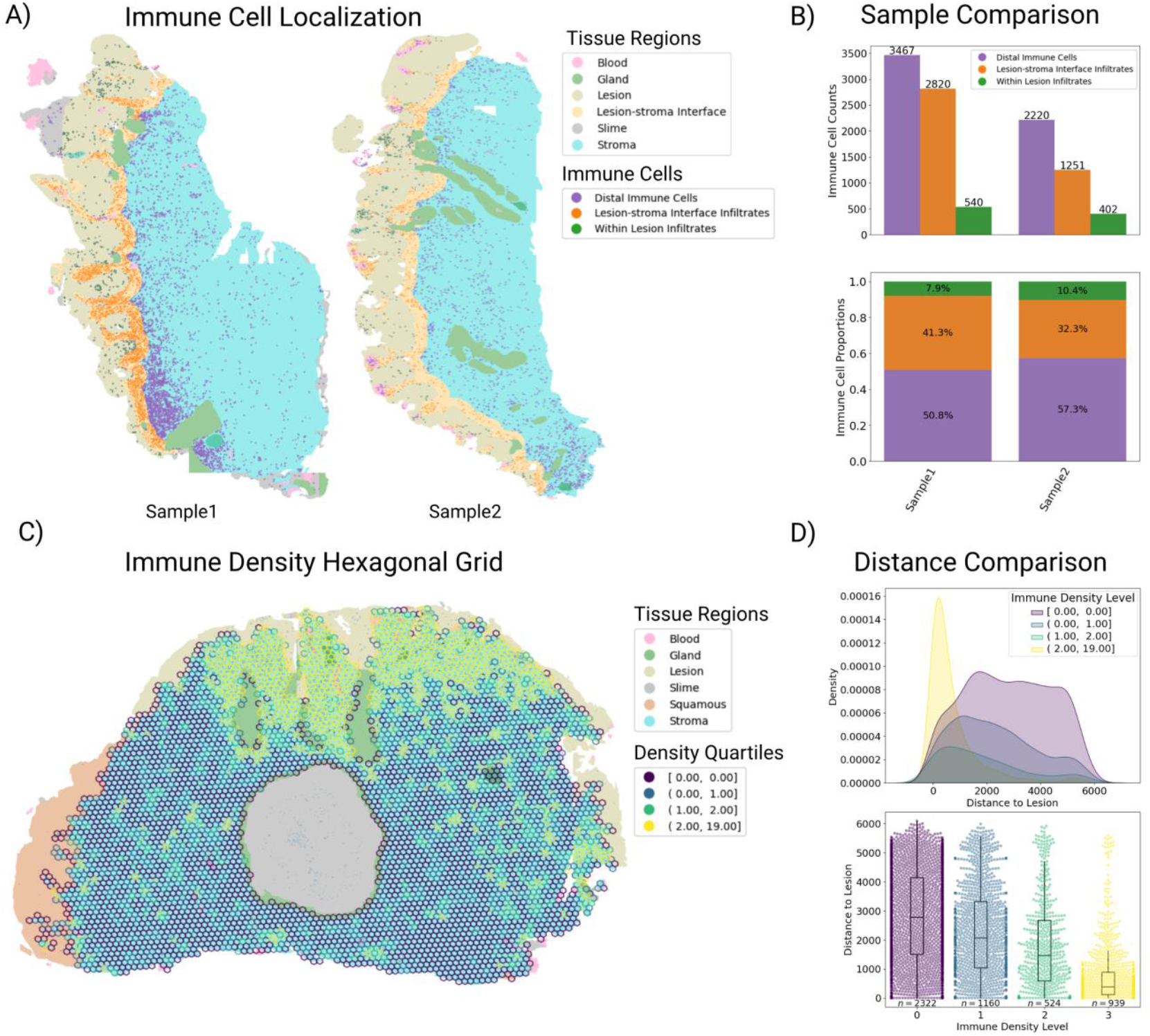
Region-based Spatial Queries of Immune Cells and Stromal Partitioning in Segmented Cervical Biopsy WSIs. **A)** Panoptic segmentation of two cervical biopsies, highlighting tissue types and immune cells categorized by their spatial location: within the neoplastic lesion, at the lesion-stroma interface, and within the distal stroma, demonstrating Histolytics’ application for partitioning and spatially querying the results of panoptic segmentation. **B)** Comparison of immune cell distributions across the two samples (absolute counts and proportional differences). In the first biopsy, 49.2% of all immune cells are located within or adjacent to lesion with especially high concentrations of immune cells located at the lesion-stroma-interface (41.3%). In contrast, in the second biopsy we see less overall infiltration as 42.7% of the immune cells are located within or adjacent to lesion with 32.3% located at the lesion-stroma-interface **C)** Application of hexagonal spatial indexing (H3) to a segmented cervical biopsy. Immune cell counts are computed per hexagon and binned into four quartiles. **D)** The observed closer localization of immune-rich hexagons to the lesion, based on Euclidean distance, suggests an activated immune response concentrated around the lesion.

To further demonstrate how Histolytics can be used to quantify spatial patterns of immune cell distributions that broad region-based queries might miss, we use Histolytics’ to partition segmented cervical stromal tissue into smaller, uniform units. By using the H3 hierarchical geospatial indexing system (https://h3geo.org/), supported by Histolytics, we subdivided the stromal compartments into hexagonal units and quantified the immune densities within each hexagon (Fig 2c). By computing the shortest distance from the center of each hexagon to the boundary of the lesion, we could quantitatively assess the spatial distribution of immune dense regions relative to the lesion. In this example, the immune-dense portions of the stroma localize significantly closer to the lesion (Fig 2d), suggesting an activated immune response.

Taken together, these examples demonstrate how Histolytics enables the analysis of the organization of segmented objects across tissues. Histolytics’ scalable spatial querying and partitioning allow the spatial organization analysis at WSI-scale without the need to use patch-level approaches.

### Morphological Profiling of Nuclei and Stroma Across WSIs

The study of morphological characteristics of nuclei and the ECM is crucial in histopathology for diagnostics, tumor grading, and predicting disease behavior^35,36^. For instance, changes in nuclear shape and chromatin patterns are well-established indicators of malignancy^36^. Similarly, the architecture and remodeling of the ECM play a significant role in tumor progression^37^.

Quantifying these types of morphological features at a large scale with high spatial resolution can provide valuable insights into disease mechanisms and heterogeneity in complicated tissue environments. Histolytics offers over 20 shape quantification metrics and several pixel-intensity-based metrics for nuclear, tissue, and ECM morphology characterization (Supplementary Table 2).

To demonstrate Histolytics’ usability for WSI-scale nuclear morphology quantification, we computed the proportion of nuclear area occupied by chromatin for every neoplastic nucleus in two ovarian high grade serous carcinoma (HGSC) slides previously segmented using Histolytics’ HGSC panoptic segmentation model. First, we partitioned the segmented tumor regions using a rectangular grid, followed by the segmentation of the chromatin clumps and mean-aggregation of the chromatin-clump-to-nucleus proportions inside each quad. This allowed us to efficiently characterize the WSI-level differences in chromatin patterns between the two slides (Fig. 3a).

**Figure 3:**
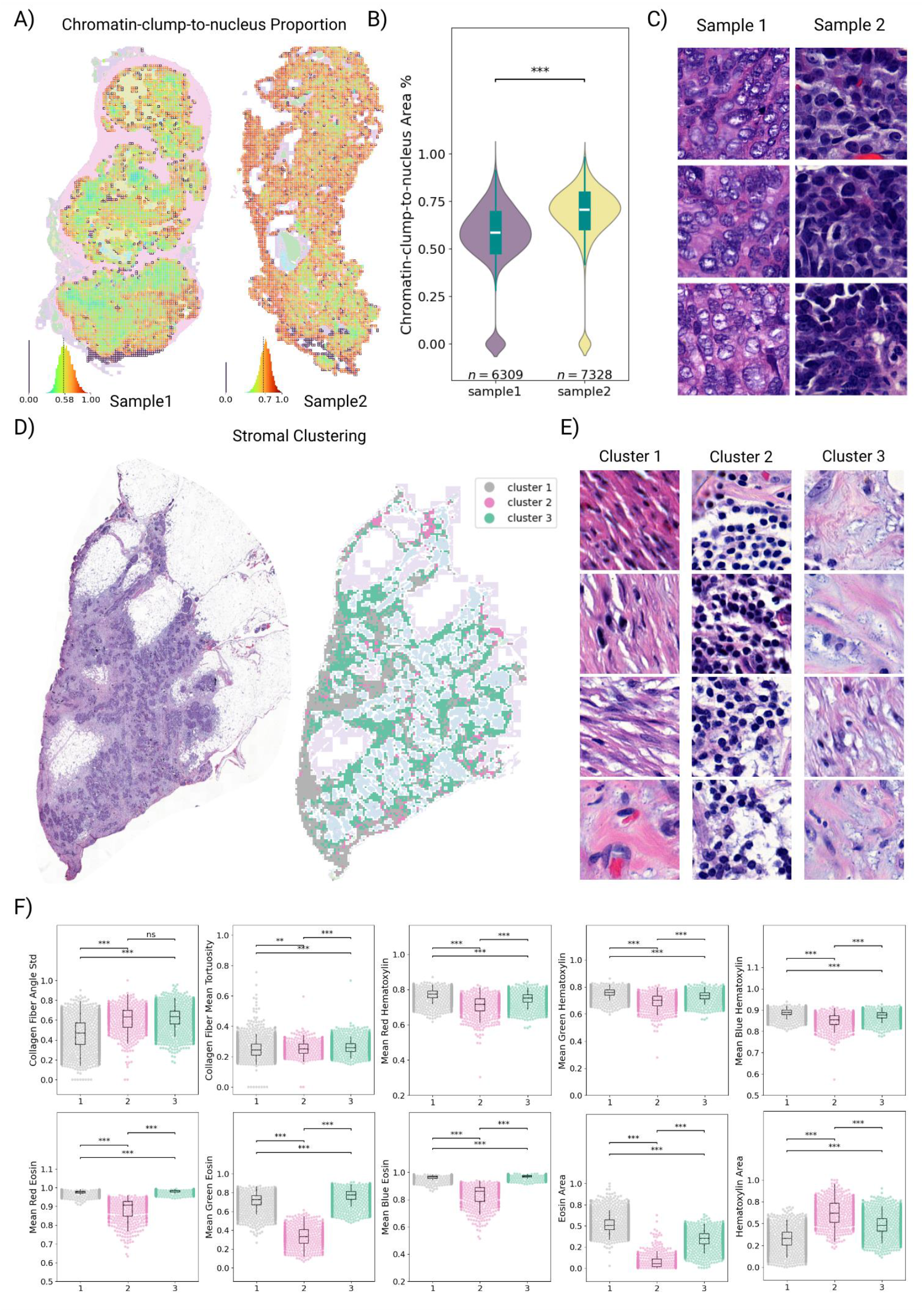
Quantification of Morphological Features in HGSC Omental Tumor WSIs. **A)** Nuclear morphological feature quantification: Panoptic segmentation of two HGSC omental slides, with a rectangular grid overlay. Within each grid-cell, neoplastic nuclei are identified, and chromatin clumps are segmented. This is followed by the computation of the mean chromatin-clump-to-nuclear area proportion, quantifying the level of chromatin distribution within neoplastic nuclei **B)** Violin plots comparing the distribution of mean chromatin-clump-to-nuclear proportions between the two samples, showing statistically significant differences determined by two-sided Mann–Whitney–Wilcoxon test **C)** Close-up examples of nuclei with chromatin-clump-to-nuclear proportions near the sample mean, illustrating the contrast between optically clear and dark hyperchromatinated nuclei. **D)** Using 10 ECM-derived morphological features computed within quads located solely within the stromal compartment of a segmented HGSC slide, we applied K-means clustering (k=3, silhouette score optimized) to dissect the stroma into its desmoplastic components. The features included the standard deviation of collagen fiber orientation (angle), mean tortuosity of collagen fibers, mean RGB values of eosin (three features), mean RGB values of hematoxylin (3 features), area of eosin component, and area of hematoxylin component. **E)** Close-up examples of stromal clusters indicative of different maturity levels of reactive stromal desmoplasia and immune density. Cluster 1 represents mature desmoplastic stroma, Cluster 2 represents immune dense stroma, and Cluster 3 represents immature desmoplastic stroma. **F)** Cluster-wise swarm plots illustrating the min-max normalized stromal features used for clustering, revealing statistically significant differences between clusters as determined by Mann–Whitney tests. Specifically, Cluster 1 is characterized by enrichment in a high eosin stain component and lower angular deviation of collagen fibers compared to other clusters. Cluster 2, in contrast, shows enrichment with a high hematoxylin stain component and a low eosin stain component, while Cluster 3 is defined by moderate levels of both hematoxylin and eosin components alongside a high angular deviation of collagen fibers. P-value annotation legend is the following: ^***^ p < 0.001, ^**^ p < 0.01, ^*^ p < 0.05, ns p > 0.05

The first slide in this example shows a broad distribution of optically clear nuclei where the chromatin is located near the nuclear membrane, while the second slide exhibits a wide-scale presence of uniformly hyperchromatic nuclei (Fig. 3b, c), illustrating distinct chromatin staining patterns associated with the metric and varying disease conditions.

To illustrate how Histolytics’ can also be used to characterize the morphology and texture of the ECM, we used it to categorize desmoplastic stromal reaction in an omental HGSC slide. Stromal reactions to cancer have variability in their textural characteristics that reflect different stages of stromal maturation and have shown to impact disease progression and therapeutic response^38^. To characterize these maturation levels within the HGSC slide, we used Histolytics to cluster the segmented stroma into three distinct compartments based on ten quantitative morphological and textural features (Fig 3d). This clustering-based analysis allowed us to dissect the stroma with histopathologic interpretability: Mature fibrotic desmoplasia, immune cell enriched stroma, and immature desmoplastic stroma were represented by clusters 1, 2, and 3, respectively (Fig 3e), with clear cluster-level feature differences (Fig 3d).

In sum, Histolytics’ comprehensive suite of nuclear and ECM morphology characterization tools showcase its ability to interpretably quantify morphological and textural heterogeneity at the WSI-level.

### Cell and ECM Organization Through Graph-based Neighborhood Analysis

The analysis of spatial relationships between different cellular and ECM components is critical for understanding complex biological processes. In cancer, for example, the interplay between tumor cells and immune cells, as well as the influence of stromal cells like cancer-associated fibroblasts (CAFs) have been shown to impact therapeutic response through immune evasion^37^. Similarly, the remodeled spatial arrangement of collagens in the ECM can influence cell migration affecting progression in many cancers^39–41^. Spatial graphs provide a powerful framework for defining proximity-based relationships and establishing spatial neighborhoods for segmented objects. Histolytics supports several spatial graph algorithms (Supplementary Table 2) to define spatial neighborhoods, such as Delaunay triangulation, Distband graphs, and K-nearest neighbor graphs (Fig. 4a). Histolytics also includes commonly used Ripley statistics^42^, and global/local spatial autocorrelation (Moran’s I^43^) tools to study spatial point patterns and clustering. It also supports custom link-based metrics, enabling flexible analysis of proximity-based neighborhoods.

**Figure 4:**
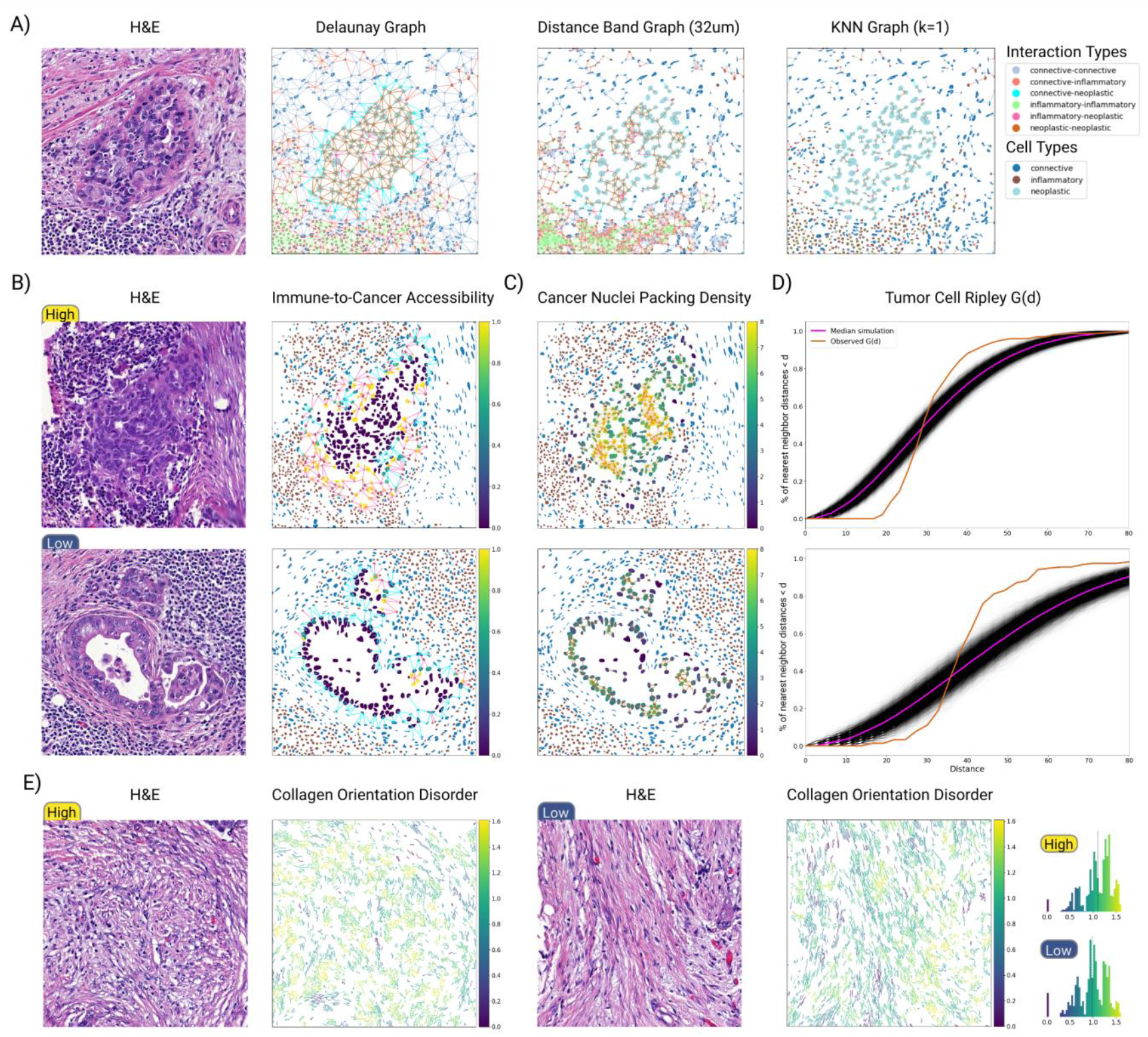
Extraction and Quantification of Graph-based Neighborhood Features in HGSC Omental Tumor Images. **A)** Examples of graph algorithms included in Histolytics, namely, Delaunay, Distband (32 m), and KNN (K=1). **B)** Immune-to-Cancer accessibility score, a metric derived by computing the proportion of immune cells to connective cells for each neoplastic nuclei neighborhood (Delaunay graph). In the first HGSC example patch, neoplastic nuclei at the tumor-stroma-interface have visibly more immune-to-neoplastic links compared to the second example, where connective cells effectively block the access of immune cells to neoplastic cells. **C)** Neoplastic nuclei neighborhood (Distband 32 m) density, measuring the number of neoplastic neighbors for every neoplastic nucleus. In the first example, the tumor cell nest shows a high-density solid growth pattern whereas in the second example, the tumor nest shows a less dense adenopapillary-like growth pattern. **D)** Ripley G-function for the neoplastic cell centroids: the cumulative distribution of nearest neoplastic neighbor distances measured over increasing distance thresholds compared to a set of simulated spatially random poisson processes. In the first example, the function increases rapidly with distance, indicating a clustered pattern whereas in the second example the increase starts later, indicating a more dispersed pattern. **E)** Example of collagen fiber orientation disorder, defined by Shannon entropy. Orientation disorder is computed for every collagen fiber at the neighborhood-level (Distband 64 m) and binned into 5 bins using the Fisher Jenks algorithm. The first example patch with high disorder exhibits more chaotic and randomly oriented collagen fibers whereas the low diversity example shows longer and mostly aligned collagen fibers.

To showcase Histolytics’ utility for graph-based neighborhood analysis, we used Histolytics to compute a potential surrogate marker for immune evasion for HGSC tumor nests by measuring the level of stromal cells physically blocking the access of immune cells to tumor. For this, we developed a simple single-cell-level metric: immune-to-cancer accessibility score, for which we computed the proportion of immune cells relative to stromal cells within each neoplastic neighborhood. To define the neoplastic neighborhoods, we used a Delaunay graph with a maximum link distance of 50 m. In this example, the tumor cell nests show clear differences in immune accessibility score (Fig 4b, Supplementary Fig 1 shows group-level comparisons at cohort level), with the second tumor nest exhibiting more extensive stromal cell barriers shielding the neoplastic cells.

As a further demonstration of Histolytics’ graph-based neighborhood analyses, we quantified the nuclear packing density in two HGSC tumor nests. To quantify this spatial metric related to tumor growth patterns we first defined the neoplastic cell neighborhoods using a Distband graph (32 m radius) and then determined the neoplastic cardinality of the neighborhoods. As expected, the solid tumor nest showed higher packing density than the adeno-papillary nest (Fig. 4c). We also measured the level of the neoplastic nuclei clustering using Ripley’s G-function over a range of distance thresholds (Fig 4d). The more rapid increase of the G-function for the solid growth pattern, indicative of a dense clustered spatial distribution, verified the same observation.

Histolytics can also be used for the graph-based neighborhood analysis of the ECM. As an example, we use Histolytics to investigate the organization of the extracellular matrix by quantifying collagen fiber orientation disorder, a prognostic factor within the tumor microenvironment known to influence cell migration and invasion^44^. For this measurement, we first used Histolytics to segment the collagen fibers from the ECM and defined collagen neighborhoods with a Distband graph (64 m radius). After this, we computed the neighborhood angular diversities using Shannon entropy^45^, representing the orientation disorder metric. In the example, the high collagen disorder sample exemplifies a more chaotic collagen neighborhood, while the low disorder sample exemplifies more aligned collagen fibers (Fig. 4e).

Collectively, these examples demonstrate that Histolytics delivers a flexible platform for in-depth neighborhood analysis, encompassing a variety of graph methods to define neighborhoods, and extensive neighborhood analysis tools for diverse use-cases.

### Analysis of Immune Cell Clusters Across WSIs

The spatial organization of immune cells within the tumor microenvironment, particularly the formation of lymphoid aggregates (Lagg) like tertiary lymphoid structures (TLS), is increasingly recognized as a critical factor influencing anti-tumor immunity and patient prognosis across various cancers, by serving as sites for immune cell activation and differentiation, impacting the local and systemic immune response to the tumor^46^. Histolytics supports multiple spatial clustering algorithms and cluster measures for cluster feature analysis (Supplementary Table 2) To investigate the spatial clustering of immune cells in HGSC, we leveraged Histolytics to perform WSI-scale cluster analysis on two H&E-stained HGSC samples. We employed density-based spatial clustering algorithm DBSCAN, supported by Histolytics, to identify cellular clusters across the entire WSIs (Fig. 5a). Immune clusters representing different sized immune cell concentrates and lymphoid aggregates such as TLS (Fig 5b) were detected from both samples. To quantify more specific cluster characteristics, we computed the total area occupied by each cluster, the number of cells contained within it, a centrography measure of the spatial spread of the cluster (dispersion), and the distance from each cluster to the nearest segmented tumor region (Fig 5c). The first example exhibited, on average, larger and more dispersed immune cell clusters, possibly indicating a larger number of activated lymphoid structures.

**Figure 5:**
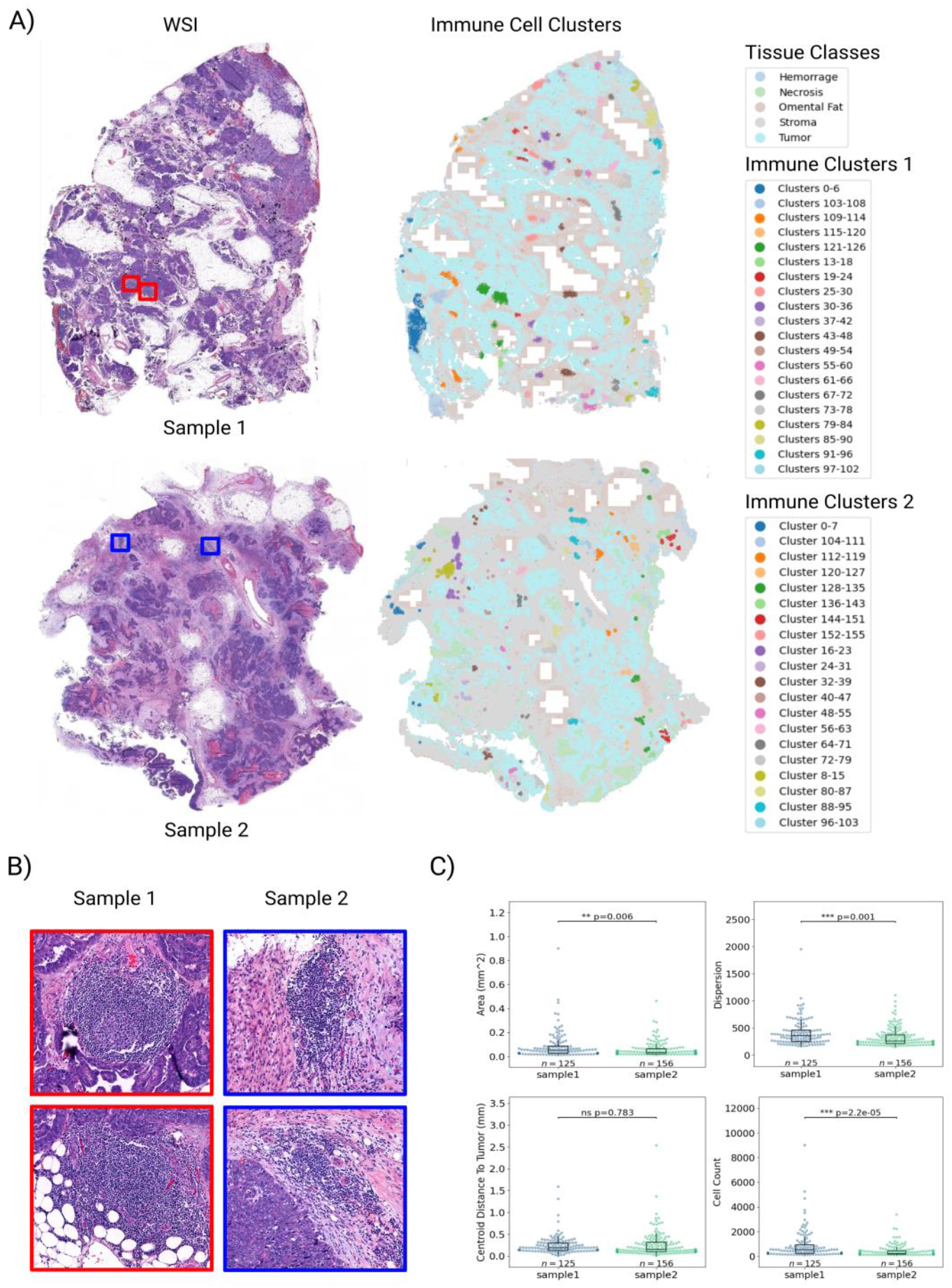
Spatial Clustering of Immune Cells and Quantification of Cluster-level Features in HGSC Omental Tumor WSIs. **A)** Two HGSC omental tumor slides and their corresponding tissue segmentation maps with spatially clustered immune cell aggregations highlighted. The spatial immune cell clusters were found using the DBSCAN algorithm (maximum distance= m, minimum size=100). **B)** Sample-wise close-up examples of Lymphoid aggregates (Lagg) and Tertiary Lymphoid Structures (TLS) which can be detected through spatial clustering. **C)** Sample-level comparison of immune cell cluster features. We computed the area, cell count, dispersion (spread), and centroid distance to tumor for each immune cluster in both samples. The first example shows significantly larger immune clusters based on higher area, dispersion and cell-counts in comparison to the second example as determined by Mann–Whitney tests.

In conclusion, the versatile, quantitative analysis capabilities of spatial cellular clusters in Histolytics’, enable the systematic exploration of cellular arrangements and clustering patterns in tissue microenvironments.

## Discussion

Histolytics introduces an efficient and scalable framework for comprehensive, interpretable analysis of H&E-stained whole-slide images (WSIs), enabling processing hundreds of slides efficiently. By tightly integrating panoptic segmentation with spatial analysis tools, such as morphological quantification, spatial querying, graph-based neighborhood analysis, and clustering, Histolytics enables consistent extraction of biologically meaningful, spatially contextualized features across diverse tissue types. The framework is designed to translate H&E images to knowledge, providing insights that are both quantitatively robust and human interpretable. With its modular design, state-of-the-art segmentation models, and seamless integration into the Python data science ecosystem, Histolytics lowers the barrier to large-scale spatial tissue analysis. Extensive documentation, tutorials, and pretrained models further support wide adoption, making Histolytics a practical tool for advancing transparent, scalable computational pathology.

## Methods

### WSI I/O

#### SlideReader

The SlideReader object, adapted and extended from the HistoPrep library (https://github.com/jopo666/HistoPrep), provides a unified interface for accessing WSIs stored on disk. It supports a diverse range of slide vendors and file formats (Supplementary Table 2) through its integration with multiple backend libraries: cuCIM^18^, OpenSlide^19^, and BioIO^20^. This multi-backend design ensures broad compatibility and leverages the strengths of each library, such as cuCIM’s efficient GPU acceleration for TIFF and SVS formats and OpenSlide’s extensive format support extended by BioIO. The SlideReader object is initialized with the path to the input WSI and the backend argument, while a region of a WSI is read from disk with XYWH-coordinates and the pyramid level arguments.

~~~
reader = SlideReader(path=PATH, backend=<str>)
tile = reader.read_region(xywh=[<int>, <int>, <int>, <int>], level=0)
~~~

Beyond basic loading, the SlideReader incorporates functionalities such as tissue detection with Otsu’s thresholding^47^ to mask out background areas and connected components-based grouping of the tiles for selecting specific regions from multi-sample slides.

~~~
thresh, tissue_mask = reader.get_tissue_mask(level=<int>)
coords = reader.get_tile_coordinates(width=<int>, tissue_mask=tissue_mask)
connected_tile_grids = get_sub_grids(coords)
filtered_tiles = [grid for grid in connected_tile_grids if len(grid) > 100]
~~~

### Panoptic Segmentation

#### Model Architectures

Histolytics incorporates a suite of deep learning multi-task model architectures specifically designed for panoptic segmentation of histopathological WSIs. These model architectures are built upon established and high-performing nuclei segmentation models: Cellpose, Stardist, Hover-Net and Cell-VIT^21–24^. To achieve panoptic segmentation capabilities for these models, we augmented them with an additional branch dedicated to semantic tissue segmentation. This parallel branch learns to delineate broader tissue regions concurrently with instance segmentation of individual nuclei. We provide several pre-trained models, easily accessible through our Hugging Face model pages (https://huggingface.co/histolytics-hub), with model weights loadable via a single line of code:

~~~
model = CellposePanoptic.from_pretrained(<str>)
~~~

Furthermore, Histolytics supports the integration of different model backbones, with tutorials (https://hautaniemilab.github.io/histolytics/user_guide/seg/backbones/) demonstrating how various panoptic segmentation models can be initialized with foundation model backbones.

#### WsiPanopticSegmenter

The WsiPanopticSegmenter object within Histolytics provides a streamlined workflow for performing panoptic segmentation on Whole Slide Images (WSIs). It takes as input a SlideReader object, a list of tile XYWH-coordinates, and a pre-trained panoptic segmentation model.

~~~
segmenter = WsiPanopticSegmenter(reader, model, coords, level=<int>)
segmenter.segment(SAVE_PATH)
~~~

Following the tile-based segmentation, the WSISegmenter can be used to merge the segmentation outputs from the individual tiles into a cohesive, whole-slide-level panoptic segmentation map. The output file sizes can be reduced with the `simplify_level` parameter that specifies the simplification level with the Douglas-Peucker algorithm^48^ and by the `precision` parameter that defines the number of decimal places the segmentation polygons are rounded to.

~~~
segmenter.merge(PATH, simplify_level=<int>, precision=<int>)
~~~

A tutorial (https://hautaniemilab.github.io/histolytics/user_guide/seg/panoptic_segmentation/) on how to run WSI-level panoptic segmentation and merging is provided, showcasing how users can efficiently process large histopathological images to generate comprehensive, whole-slide-level panoptic segmentation maps.

### Model Training and Benchmarking Utilities

Histolytics provides a comprehensive set of utility functions designed to streamline the training and benchmarking of panoptic segmentation models. Recognizing the complexities of training multi-task architectures, the library includes several commonly used segmentation loss functions (Supplementary Table 2) alongside a flexible `MultiTaskLoss` wrapper which enables the simultaneous optimization of the different output branches in panoptic segmentation models with customizable weighting schemes. Furthermore, Histolytics incorporates advanced regularization techniques to enhance model generalization and performance, including the implementation of the StrongAugment^49^ augmentation pipeline and several loss-weighting and regularization techniques embedded within the Loss-functions (Supplementary Table 2). We provide an extensive tutorial (https://hautaniemilab.github.io/histolytics/user_guide/seg/finetuning/) on how to train panoptic segmentation models from scratch with new data, enabling users to adapt the models to their specific datasets and research questions.

To allow for quantitative assessment of model performance across different aspects of the panoptic segmentation task, including both instance segmentation of individual nuclei and semantic segmentation of tissue regions, Histolytics provides a wide array of benchmarking metrics (Supplementary Table 2). For example, the library includes standard metrics commonly used in the field, such as Panoptic Quality (PQ), Average Precision (AP), as well as metrics for semantic segmentation performance, including class-wise Intersection over Union (IoU) and DICE coefficients.

### Spatial Analysis

#### Spatial querying

Histolytics includes an efficient spatial querying engine for segmented objects. The `get_objs` function enables rapid subsetting of segmented entities, such as individual nuclei or other cellular components, based on spatial predicates with designated regions of interest, such as segmented tissue areas or user-defined boundaries. Leveraging the fast R-tree-based spatial indexing system of geopandas^26^ library, the function accepts a set of segmented objects (geopandas.GeoDataFrames) and a target spatial region (geopandas.GeoDataFrame) and return a geopandas.GeoDataFrame containing the queried output objects.

~~~
nuclei_within_roi = get_objs(roi, segmented_objects, predicate=“contains”)
~~~

Tutorial (https://hautaniemilab.github.io/histolytics/user_guide/spatial/querying/) on spatial querying are provided, demonstrating how users can efficiently extract specific objects based on their spatial relationships in panoptic segmentation maps.

#### Spatial partitioning

Histolytics offers flexible methods for spatially partitioning tissue segmentations. The library implements buffering-based techniques to delineate interface regions between distinct tissue types. By applying a user-defined buffer around the boundary of one tissue region, overlapping areas with an adjacent tissue can be extracted and analyzed, enabling the study of interactions and transitions at tissue interfaces.

~~~
interface = get_interfaces(tissue1, tissue2, buffer=100)
~~~

Furthermore, Histolytics provides functionality to partition tissue regions into regular spatial grids, including hexagonal grids and rectangular grids. By leveraging the H3 and Quadbin hierarchical geospatial indexing systems or a regular rectangular grid-fitter, tissue regions can be subdivided into smaller, uniform units, allowing for the quantification of spatial features such as cell densities, morphological characteristics, or marker expression within each grid cell.

~~~
h3_gr = h3_grid(tissue, resolution=<int>)
quadbin_gr = quadbin_grid(tissue, resolution=<int>)
rect_gr = rect_grid(tissue, resolution=(<int>, <int>), overlap=<int>)
~~~

Tutorial (https://hautaniemilab.github.io/histolytics/user_guide/spatial/partitioning/) on spatial partitioning are also available, where we show how segmented tissue regions can be partitioned for more localized analysis

#### Chromatin Clump Segmentation

Histolytics provides `chromatin_feats` function to segment dense chromatin regions within identified nuclei (defined by a 2D label map). The segmentation algorithm is based on multi-Otsu thresholding^47^ (Scikit-image^50^ implementation) of the grayscale input image which is masked by the label map. The function returns a binary mask of chromatin clumps.

~~~
chrom_mask = chromatin_feats(img, label)
~~~

#### Nuclear Features

Histolytics includes a comprehensive collection of nuclear features with 17 shape quantification metrics, 13 intensity quantification metrics for both grayscale and RGB intensities (total of 52 features), 10 textural metrics (Scikit-image^50^ implementations), and five chromatin clump distribution metrics (Supplementary Table 2). Designed to characterize nuclear pleomorphism, these features capture a wide range of nuclear attributes, from basic geometry and size to complex textural and intensity patterns.

These metrics can be computed using the `shape_metric`, `grayscale_intensity_feats`, `rgb_intensity_feats`, `chromatin_feats`, and `textural_feats` functions. The `shape_metric` takes segmented objects as input geopandas.GeoDataFrame and returns the input with the metrics added as new columns. The rest of the functions take in an H&E image and it’s corresponding nuclei label mask in raster format and return pandas.DataFrame objects containing the computed features per nuclei.

~~~
morphometrics = shape_metric(nuclei_gdf, metrics=[<str>])
gray_feats = grayscale_intensity_feats(img, label_mask, metrics=[<str>])
rgb_feats = rgb_intensity_feats(img, label_mask, metrics=[<str>])
chrom_feats = chromatin_feats(img, label_mask, metrics=[<str>])
texture_feats = textural_feats(img, label_mask, metrics=[<str>])
~~~

We provide an extensive tutorial (https://hautaniemilab.github.io/histolytics/user_guide/spatial/nuclear_features/) on nuclear feature extraction, enabling users to quantify a wide array of nuclear-level features from images and nuclei segmentation maps.

#### Collagen Fiber Segmentation

Histolytics includes `extract_collagen_fibers` function to segment collagen fibers within histological images. The function is based on the Scikit-image^50^ implementation of the Canny edge detection algorithm^51^ that is applied to the grayscale version of the input image to identify potential fiber structures. To refine the collagen fiber segmentation, edges are detected only within the identified stromal parts of the image by masking out high eosin stained and background regions from the image.

~~~
collagen_mask = extract_collagen_fibers(img)
~~~

#### Stromal Features

Histolytics enables the extraction of eight features related to the structural organization of extracted collagen fibers with the `fiber_feats` function. The computed features are designed to measure the length, orientation, curviness and tortuosity of the collagen fibers (Supplementary Table 2).

~~~
collagen_feats = fiber_feats(img, metrics=[<str>])
~~~

Furthermore, for stromal regions of the H&E images, the `stromal_intensity_feats` function can be used to compute intensity features of the stroma. The function first decomposes the RGB image into Hematoxylin (H), Eosin (E), and DAB (D) channels using Scikit-image^50^ implementation of the HED-color deconvolution method. Binary masks for eosinophilic and hematoxylinophilic stain components within the stroma are generated using Otsu thresholding^47^ on the respective color channels, while excluding areas occupied by cells and background. The function returns a sequence of 13 intensity metrics for each RGB channel within the segmented eosin and hematoxylin regions of the stroma (total 81 features at most), along with the area occupied by both Eosin and Hematoxylin stain (Supplementary table 2).

~~~
intensity_feats = stromal_intensity_feats(img, metrics=[<str>])
~~~

We provide an extensive tutorial (https://hautaniemilab.github.io/histolytics/user_guide/spatial/stromal_features/) showcasing how users can efficiently characterize stromal features in their H&E images.

#### Graphs

Histolytics leverages graph-based representations to quantify neighborhood relationships and spatial interactions between segmented objects. In a graph, a segmented object is a node and a link between two nodes is defined by a graph algorithm and a distance threshold (links can be formed between nodes that are spatially close to each other). A node and all its links to other nodes form a local neighborhood.

In Histolytics, the graph implementations rely on the Libpysal^25^ library and supported graph types include, but are not limited to, Delaunay triangulation, distance-based graphs (Distband), and KNN-graphs (full list in Supplementary Table 2). The `fit_graph` function serves as a wrapper for these algorithms with additional capability to filter edges based on a given distance threshold. The graph is returned as a geopandas.GeoDataFrame and a `libpysal.weights.W` object that contains methods for easy manipulation of the graph and conversion to scipy^52^ sparse adjacency matrices or networkx.Graph^53^ objects for more sophisticated network analysis.

w, w_gdf = fit_graph(segmented_objects, method=<str>, threshold=<int>)

A tutorial (https://hautaniemilab.github.io/histolytics/user_guide/spatial/graphs/) on graph fitting and graph centrality feature extraction is provided, enabling users to leverage graph based feature extraction methods easily.

#### Neighborhood Features

Histolytics provides functionalities to characterize local neighborhoods of segmented objects through the `local_character`- and `local_distances` functions. The `local_character` function enables the computation of summary statistics (e.g., mean, standard deviation, sum) for any previously extracted feature within the neighborhood of each segmented object (full list in Supplementary table 2). This allows for the summarization of local neighborhoods based on the distribution of specific features in the vicinity of each nucleus or other segmented entity. In contrast, the `local_distances` function focuses specifically on measuring the Euclidean distances between neighboring segmented objects. This function calculates the pairwise distances between nodes within a defined local neighborhood, enabling the analysis of object spacing and packing arrangements. Both functions take in as input the spatial graph of the segmented objects, the geopandas.GeoDataFrame of the segmentations and returns the input geopandas.GeoDataFrame with added columns for each computed metric.

objs = local_character(w, objs, reductions=[<str>]) objs = local_distances(w, objs, reductions=[<str>])

Furthermore, Histolytics enables the quantification of local neighborhood heterogeneity of segmented objects using a variety of diversity indices. Computed at the object level (e.g., nuclei, collagen), these indices characterize the diversity of the immediate surroundings of each segmented entity. The `local_diversity` function provides a wrapper for computing metrics such as the Simpson Diversity Index^54^, Shannon Entropy^45^, Gini Index^55^, and Theil Index^56^. These measures assess the compositional heterogeneity within each objects neighborhood, allowing for the analysis of spatial diversity patterns across the tissue microenvironment and their potential association with other spatial features. The `local_diversity` function takes in the spatial graph of the segmented objects, the geopandas.GeoDataFrame of the segmentations and a list of the specified diversity metrics and returns the input geopandas.GeoDataFrame with added columns for each computed diversity metric.

objs = local_diversity(w, objs, metrics=[<str>])

A comprehensive tutorial (https://hautaniemilab.github.io/histolytics/user_guide/spatial/nhoods/) showcasing the usage of neighborhood feature extraction is provided.

#### Ripley Statistics

Ripley functions^42^ are a group of spatial statistics that focus on how to characterize clustering in point patterns through pairwise and nearest neighbor distances. Ripley functions characterize point patterns as a function of distance with the aim of determining whether the point patterns are random, dispersed or clustered. Histolytics contains implementations of Ripley’s K, L and G functions.

Ripley’s K-function estimates the expected number of points within a certain distance of a randomly chosen point. By calculating this across a range of distances, the K-function returns a vector of cumulative average number of points lying within a given distance of a typical data point. The Ripley’s L-function is a transformation of the K-function that stabilizes the variance of the K-estimator and transforms it into a straight line making the visual assessment of the function easier.

The Ripley’s *G-*function, on the other hand, summarizes the distribution of the nearest neighbor distances between each point in a point pattern. This is done by computing the proportion of points for which the nearest neighbor is within a given distance for a range of distance thresholds. Ripley’s G-function returns a vector of cumulative percentages against increasing distances.

The Ripley’s functions are typically computed for empirical data and compared against a completely spatial random Poisson process. In Histolytics, this can be done using the `ripley_test` function that takes in as an input segmented objects and a range of distances and returns the computed Ripley’s function distribution, the random simulated spatial process distribution of the Ripley’s function, and the results of the statistical tests against the random process.

ripl, sim, pval = ripley_test(objs, distances=[<int>], type=<str>)

#### Global and Local Spatial Autocorrelation

Spatial autocorrelation refers to the degree to which features or values are correlated to each other across space. Positive spatial autocorrelation indicates that nearby objects tend to have similar values, while negative spatial autocorrelation suggests that nearby objects tend to have dissimilar values. To quantify spatial autocorrelation, Histolytics implements Moran’s I statistic which can be used to assess whether the distribution of a specific feature’s values across the segmented objects is spatially random, clustered, or dispersed.

Histolytics has implementations for global and local Moran’s I statistics via the `global_autocorr` and `local_autocorr` functions, respectively. These functions serve as wrappers for Moran’s I implementations within the Esda module of the Libpysal library^25^. The functions take as input a geopandas.GeoDataFrame of segmented objects and a spatial graph representing the neighborhood relationships between these objects. The `global_autocorr` function returns a single global Moran’s I value and its associated p-value across all segmented objects, measuring the significance of autocorrelation against a random spatial process of a given feature.

Conversely, the `local_autocorr` function computes the local Moran’s I statistic and its corresponding p-value for each individual segmented object within the input geopandas.GeoDataFrame returning arrays of Moran’s I statistics and p-values.

~~~
I, p_sim = global_autocorr(segmented_objs, G)
Il, p_sim, quadrants = local_autocorr(segmented_objs, G)
~~~

#### Spatial Clustering and Centrography

Histolytics provides two distinct approaches for identifying spatial clusters of segmented objects. First, it implements density-based clustering methods (Supplementary Table 2), primarily leveraging the scikit-learn^57^ implementations of DBSCAN algorithm^58^ and its variants. These clustering algorithms are accessed via `density_clustering` function wrapper. Second, Histolytics offers clustering based on Local Indicators of Spatial Association^59^ (LISA) via the `lisa_clustering` function. This method identifies clusters of objects with similar values for a specific feature, based on the local spatial autocorrelation as measured by local Moran’s I.

~~~
labels1 = density_clustering(segmented_objs)
labels2 = lisa_clustering(segmented_objs, feature=<str>)
~~~

The resulting clusters, whether derived from density or LISA-based methods, can be further analyzed using point pattern measures (cluster size, area, orientation, dispersion) and distance measures to specified tissues (Supplementary Table 2) with the `cluster_feats` and `cluster_dists_to_tissue` functions.

~~~
cluster_feats = cluster_feats(labels)
cluster_dists = cluster_dists_to_tissue(labels, tissues, tissue_class=<str>)
~~~

We provide a tutorial (https://hautaniemilab.github.io/histolytics/user_guide/spatial/clustering/) showcasing the easy usage of density based clustering and cluster feature extraction methods.

## Data Availability

To facilitate the reproducibility of the analyses presented in this manuscript, processed datasets are made available through the histolytics.data module. The provided segmentation and image data are sufficient to fully replicate and demonstrate the analytical workflows described herein. The original image data from which these segmentations were derived originated from the Helsinki University Hospital (HUS), and the DECIDER cohort (Multi-layer Data to Improve Diagnosis, Predict Therapy Resistance and Suggest Targeted Therapies in HGSOC; ClinicalTrials.gov identifier: NCT04846933). Details regarding the preprocessing and panoptic segmentation of the original image data can be found within the segmentation pages of the Histolytics user guide documentation https://hautaniemilab.github.io/histolytics/user_guide/.

## Code Availability

Histolytics is distributed as a pip-installable Python package, and the source code is accessible in the GitHub repository: https://github.com/hautaniemilab/histolytics. Comprehensive user guides and API reference are provided in the accompanying documentation: https://hautaniemilab.github.io/histolytics/. The analytical workflows necessary to reproduce the findings presented in this manuscript are integrated within the WSI Analysis Workflows section of the User Guide documentation pages https://hautaniemilab.github.io/histolytics/user_guide/.

## Acknowledgements

We acknowledge CSC-IT Center for Science, Finland, for computational resources, Kari Lavikka for his invaluable assistance in the creation and refinement of the figures presented in this manuscript. This work utilized High-Grade Serous Carcinoma (HGSC) data, which originated from the DECIDER cohort (Multi-layer Data to Improve Diagnosis, Predict Therapy Resistance and Suggest Targeted Therapies in HGSOC; ClinicalTrials.gov identifier: NCT04846933) that has received funding from the European Union’s Horizon 2020 research and innovation programme under grant agreement No 965193. Additionally, we acknowledge Helsinki and Auria Biobanks for scanning the whole-slide data used in this work and the support from Finnish Medical Foundation. O.L acknowledges Finnish Society for Colposcopy r.y for a personal research grant.

---

**Supplementary Figure 1.**
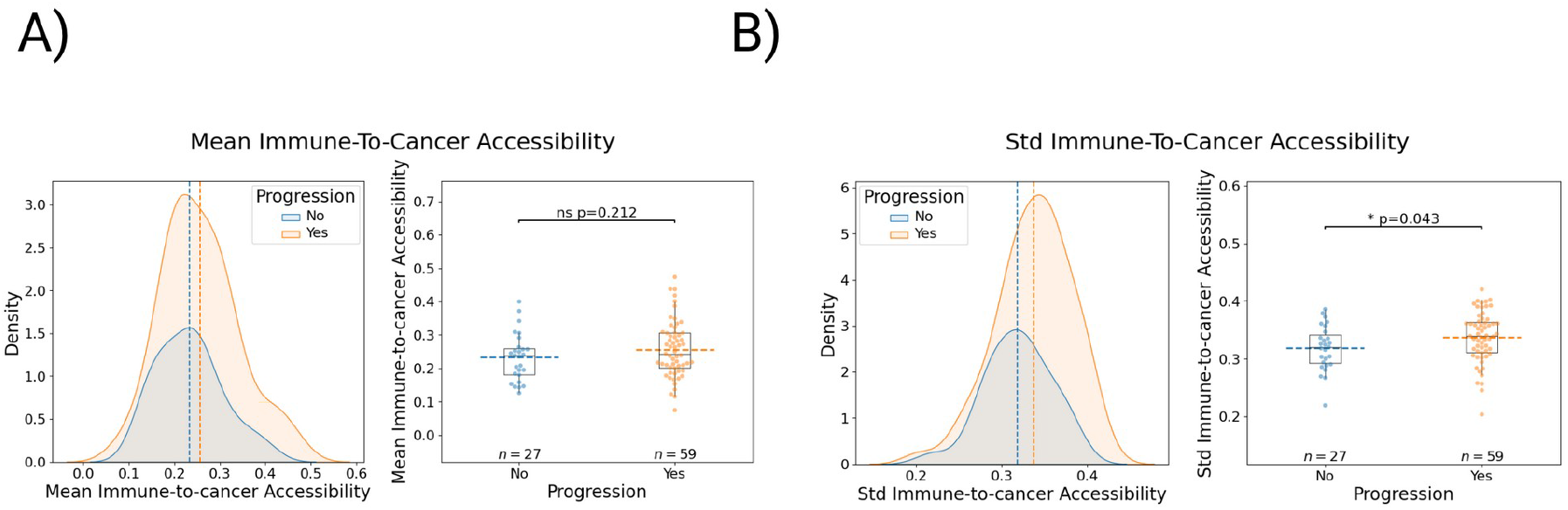
Cohort level comparison of immune-accessibility metrics in HGSC Omental Tumor Samples. **A)** The mean Immune-to-cancer accessibility score between progression free and progressing HGSC patients. No significant difference (Mann–Whitney U-test) is observed at patient-level (total 368 WSIs from 86 patients) **B)** the cell-cell variability (standard deviation) at patient level in immune-to-cancer accessibility is significantly higher (Mann-Whitney U-test) in the progressing group, indicating greater heterogeneity of how easily immune cells can reach cancer cells.

